# Micronutrient composition and microbial community analysis across diverse landraces of the Ethiopian orphan crop enset

**DOI:** 10.1101/834564

**Authors:** Solomon Tamrat, James S. Borrell, Manosh K. Biswas, Dawd Gashu, Tigist Wondimu, Carlos A. Vásquez-Londoño, Pat J.S. Heslop-Harrison, Sebsebe Demissew, Paul Wilkin, Melanie-Jayne R. Howes

**Affiliations:** Department of Plant Biology and Biodiversity Management, Addis Ababa University, Addis Ababa, Ethiopia; Department of Biology, Dilla University, SNNPR, Ethiopia; Royal Botanic Gardens, Kew, Richmond, Surrey, TW9 3AE, UK; Department of Genetics and Genome Biology, University of Leicester, LE1 7RH, UK; Center for Food Science and Nutrition, Addis Ababa University, Addis Ababa, Ethiopia; Faculty of Sciences, Universidad Nacional de Colombia, Bogotá, Zip Code 111321, Colombia; Gullele Botanic Garden, P. O. Box 153/1029, Addis Ababa Ethiopia

**Keywords:** *Ensete ventricosum*, Ethiopia, fermentation, free amino acids, food security, kocho, micronutrients, orphan crops

## Abstract

Enset (*Ensete ventricosum*) is a major starch staple and food security crop for 20 million people. Despite substantial diversity in morphology, genetics, agronomy and utilization across its range, nutritional characteristics have only been reported in relatively few landraces. Here, we survey nutritional composition in 22 landraces from three enset growing regions. We present mineral characterization of enset corm tissue, free amino acid characterization of raw and processed (fermented) tissues and genomic analysis of the microbial community associated with fermentation. We show that compared to regionally important tubers and cereals, enset is high in calcium, iron, potassium and zinc and low in sodium. We report changes in free amino acid composition due to processing, and establish that the bacteria genera *Acetobacter*, *Lactobacillus* and *Bifidobacterium*, predominate during fermentation. Nutritional and microbial variation presents opportunities to select for improved composition, quality or safety with potentially significant impacts in food security and public health.

## 1. Introduction

Humans currently satisfy most dietary requirements with surprisingly few species (Borrell et al., 2019), yet a much greater diversity of nutritionally suitable plants have been identified, often with narrow regions of utilization (Mayes et al., 2012). The food products of the large perennial herb *Ensete ventricosum* (Welw.) Cheesman (Musaceae) are an important dietary starch source in Ethiopia (Borrell et al., 2018; Fanta & Satheesh, 2019; Negash & Niehof, 2004). Commonly known as enset (or alternatively as the false banana or Abyssinian banana), this major food crop is principally cultivated as a highly resilient staple that withstands a wide range of environmental conditions and can buffer seasonal variation in food availability. Enset contributes to the food security of over 20 million people, but is virtually unknown outside of its narrow zone of cultivation in South West Ethiopia, despite growing undomesticated and unutilized across much of East and Southern Africa (Borrell et al., 2018). In addition to being processed for multiple food products, enset is also used for livestock fodder, packaging materials, fiber and traditional medicine (Borrell et al., *in press*; Mohammed, Martin, & Laila, 2013; Olango, Tesfaye, Catellani, & Pè, 2014).

Despite the importance of enset for food security and the existence of hundreds of diverse landraces (Borrell et al., 2018), the nutritional composition of the raw plant tissues and processed food products (e.g. *kocho*) has only been reported in a small number of landraces (Bosha et al., 2016; Daba & Shigeta, 2016; Mohammed et al., 2013; Nurfeta, Tolera, Eik, & Sundstøl, 2008). Reported nutritional composition and the relative concentrations of certain micronutrients show considerable variation. Whilst this may be attributable in part to differing analysis methods, the extensive diversity of genetically differentiated enset landraces (Borrell et al., 2018; Tobiaw & Bekele, 2011), heterogeneity of farm management practices (Garedew, Ayiza, Haile, & Kasaye, 2017; Olango et al., 2014) and environmental conditions, particularly soil (Amede & Diro, 2005; Borrell et al., 2018), may also be a contributing factors. Therefore, we highlight the need to profile micronutrient composition across a representative subset of enset landrace diversity, whilst also characterizing the ubiquitous effects of fermentation on the composition, relevant to the quality and safety of enset foods, and the associated microorganism diversity responsible for mediating tissue processing.

The edible parts of enset comprise the starch rich pseudopetioles forming the pseudostem (overlapping leaf sheaths) which are decorticated, and the corm (the underground base of the stem that serves as a storage organ) which is pulverized and pressed (Borrell et al., 2018). These two main tissues are collectively processed, using fermentation pits, into starch staples including *bulla* and *kocho* (see Birmeta, Bakeeva, & Passoth, 2018, for a detailed description). *Kocho* is the bulk of the fermented product and is baked into a thin fibrous bread considered to have a good shelf life. *Bulla* is a small amount of water-insoluble starchy product separated from the *kocho* during processing by squeezing and sometimes consumed separately. The corm of enset is also occasionally consumed boiled, much like potato, and this is called *amicho*.

The precise fermentation practice is variable among regions and cultural groups (Garedew et al., 2017; Hunduma & Ashenafi, 2011). Karssa et al., (2014) reports preparation of a starter culture (known as *gamancho* or *gamma*) from selected corms of mature plants, followed by a two-phase process, with surface and then pit fermentation. Bosha et al. (2016) reports a ground mix of several other plant species being added to the mashed corm to initiate fermentation. Whereas many other authors (Birmeta et al., 2018; Gashe, 1987), indicate that fermentation is initiated simply from mashed tissue left for several days at ambient temperature.

The microorganisms responsible for fermentation alter the chemical composition of the raw substrate, which in some cases enriches the nutritional value of fermented products (Tamang, Watanabe, & Holzapfel, 2016) by removing anti-nutritionals and breaking down complex components. Furthermore, microbial communities introduced during processing are often critical to food safety and preservation by preventing growth of spoilage and toxic organisms. These microorganisms, often occurring as communities in food products are poorly known in orphan and minor tropical crops cultivated by subsistence farmers (Tamang et al., 2016). However, improvement of these cultures represents a relatively accessible opportunity to enhance the nutritional consistency, bioavailability and quality of neglected food products (Chelule, Mokoena, & Gqaleni, 2010), while urbanization and penetration of shops leads to loss of cultures traditionally maintained by smallholders.

In a study on *kocho* production from enset, Gashe (1987) reported that *Leuconostoc mensenteroides* and *Streptococcus faecalis* are responsible for initiating fermentation and reducing the pH. These are then superseded by the homofermentative bacteria *Lactobacillus coryniformis* subsp. *coryniformis* and *L. plantarum* which reduced the pH further. More recently, in an analysis of kocho samples from the Wolkite area, Weldemichael, Shimelis, Emire, & Alemu (2019) identified kocho-associated bacteria as predominantly from the genera *Lactobacillus* and *Acetobacteria*, whilst a survey of kocho samples from the Gamo highlands identified high relative abundance of *Enterobacteriaceae* and *L. mesenteroides subsp*. *cremoris* in the early stages of fermentation (Andeta, Vandeweyer, Woldesenbet, Eshetu, & Hailemicael, 2018). More generally, the process of fermenting enset has been reported to reduce total protein and carbohydrates, whilst increasing free amino acids (FAAs) 1.6-fold (Urga et al. 1997). However, the specific free amino acids in fermented products have not been characterized. Therefore whether enset is a source of essential amino acids (those the body cannot synthesize in sufficient quantities) in this form is unknown.

We hypothesize that significant genomic, phenotypic, environmental and agronomic variation in enset should also result in variation in nutritional composition across landraces. In this study we investigate selected micronutrients with a focus on inorganic heavy metal and trace elements and free amino acids in domesticated enset in Ethiopia. We profile both raw tissues and processed (fermented) products in samples from 22 landraces across three major enset growing regions, and present a quantitative genomic survey of the microbial community associated with enset fermentation. We place our results in the context of other regionally available staples and the opportunities that nutritional and microbial variation presents for applied biotechnology to improve the nutritional quality, consistency and safety of enset derived foods.

## 2. Material and Methods

### 2.1 Sample collection and preparation

Fresh samples of six enset tissues (corm, pseudostem, leaf, fruit flesh, fruit peel (exocarp) and seed) were harvested from a selection mature enset plants (n=28, landraces=22) occurring in three regions; Sidama, Wolaita and Gurage zones in the Southern Nations Nationalities and Peoples Region (SNNPR), Ethiopia (Table 1). In selecting samples we sought to capture the full range of phenotypic and cultural variation. Selected plants were individually prepared, processed and fermented following traditional local practices and associated qualitative indigenous knowledge related to processing was recorded. Subsequently, samples of three enset products *kocho*, *bulla* and mixed *kocho* and *bulla* (Ko-Bu; whereby the *bulla* liquid is not isolated during preparation, a common practice in parts of Sidama) from selected plants were collected post-fermentation, air dried and powdered using an electric grinder.

**Table 1.** Enset tissue and food product types evaluated in this study.

| Type | Tissue | Samples | Uses and food products |
| --- | --- | --- | --- |
| Tissue | Corm | 19 | A raw, underground storage organ used to make both kocho and bulla. Also boiled and eaten as 'Amicho'. |
|  | Pseudostem | 9 | A raw aboveground tissue comprising overlapping leaf sheaths. The pseudostem is the main tissue type making up the bulk of both kocho and bulla. Petioles and leaf midribs are fed to cattle as fodder. |
|  | Leaf | 1 | Used as a protective layer in baking bread and to line the kocho processing area and fermentation pits |
|  | Fruit flesh | 1 | Not currently utilized |
|  | Fruit peel | 1 | Not currently utilized |
|  | Seed | 2 | Not currently utilized |
| Product | Kocho | 11 | A traditional fermented flatbread |
|  | Kocho-Bulla (KoBu) | 8 | A traditional fermented flatbread, but made without extracting the bulla liquid. In this study this practice occurs in Sidama region |
|  | Bulla | 8 | The water-insoluble starchy liquid extracted from kocho by squeezing. Bulla can be dried and used as a flour, or made into a gelatinous porridge. It is produced in small amounts and often considered the most valuable enset food product. |

### 2.2 Micronutrient and mineral composition

Micronutrient content used only raw corm tissue, as relative concentration of micronutrients should not be affected by fermentation. Total ash content was estimated using 5 g of desiccated corm tissue. Samples were charred in a hot plate (Wagtech Model ST15, Sweden) and ignited at 550°C for 5 hours a in a Muffle furnace (Carbolite Model S302RR, Sweden). Residual ash was calculated as a percentage of the original desiccated sample weight.

Mineral content was assessed using 0.5 g corm samples digested in a closed high-performance microwave digestion system (ETHOS) with the addition of trace metal grade 65% HNO_3_ (Fisher Scientific, UK) and H_2_O_2_according to the manufacturer’s recommendations under high pressure and temperature. The digest was washed with distilled water (Milli-Q). Multi-elemental analysis was conducted using an inductively coupled ICP-MS (Thermo-Fisher Scientific iCAP-Q; Thermo Fisher Scientific, Bremen, Germany). Samples were introduced from an auto-sampler (Cetac ASX-520) incorporating an ASXpress™ rapid uptake module through a perfluoroalkoxy (PFA) Microflow PFA-ST nebuliser (Thermo Fisher Scientific, Bremen, Germany). Internal standards were introduced to the sample stream on a separate line via the ASXpress unit and included Ge (10 µg L-1), Rh (10 µg L-1) and Ir (5 µg L-1) in 2% trace analysis grade (Fisher Scientific, UK) HNO_3_. External multi-element calibration standards (Claritas-PPT grade CLMS-2 from SPEX Certiprep Inc., Metuchen, NJ, USA) including Al, As, Cd, Ca, Co, Cr, Cs, Cu, Fe, K, Mg, Mn, Mo, Na, Ni, P, Pb, S, Se, and Zn, in the range 0 – 100 µg L-1 (0, 20, 40, 100 µg L-1) were used. A bespoke external multi-element calibration solution (PlasmaCAL, SCP Science, France) was used to create Ca, Mg, Na and K standards in the range 0-30 mg L-1. In addition, KH_2_PO_4_ and K_2_SO4 solutions were used to calibrate the machine during determination of P and S, respectively. Sample processing was undertaken using Qtegra™ software (Thermo-Fisher Scientific) utilizing external cross-calibration between pulse-counting and analogue detector modes when required. Blank digests containing all except enset samples were used for quality control. In addition, a certified reference material (SRM 1567b – wheat flour) from the National Institute of Standards and Technology (NIST), USA was used for standardization. Comparative plant nutritional composition data was sourced from the USDA Food Composition Databases (USDA, 2012) and Feedipedia (Heuze, Thiollet H., Tran G., Hassoun P., & Lebas F., 2017), and plotted together with data from this study.

### 2.3 Crude protein and free amino acid composition

Crude protein was estimated using 0.5 g of desiccated corm tissue. Briefly, 6ml of conc H_2_SO_4_and 3.5ml of H_2_O_2_were added followed by 3 g of a catalytic mixture of CuSO_4_and K_2_SO_4_ (1:15) for 15 minutes. The solution was heated at 370°C using a kjeldahl digester until a clear solution was observed and distilled using the Kjeldahl distillation apparatus (Auto K9840 Analyzer, Kjeltec, BCR-Technology Co, Ltd., China). The auto distiller was adjusted to add 30 ml distilled water (to avoid precipitation of sulfate in the solution), and 40 ml NaOH (35%) in order to neutralize excess acid, break down ammonium sulfate and release ammonia gas. The distillate was collected for 8 minutes in a 250 ml Erlenmeyer flask containing 2% boric acid and five drops of methyl red indicator. The distillate was titrated with standardized 0.1M HCl and sample N content determined by subtracting the N content of blank control samples.

Free amino acid composition was assessed in a subset of samples using an EZ:faast^TM^ kit (Phenomenex, Macclesfield, UK). Powdered samples (100 mg) were extracted in 100% water and sonicated for 20 min prior to centrifugation. Supernatants were derivatized using the method described in the manufacturer’s instructions, and as described previously (Dziągwa-becker, Weber, Zajączkowska, & Oleszek, 2018). Analyses of derivatized amino acids were performed with 2µl sample injections on a Thermo Scientific system, consisting of an ‘Ultimate 3000’ UHPLC unit and an ‘LTQ Velos Pro’ mass spectrometer (Thermo Scientific, Waltham, MA, USA). Chromatography (qualitative and quantitative analyses) was performed as described in the EZ:^TM^ faast kit’s instructions, with amino acids assigned by comparison with the calibration amino acid standards provided with the kit. For each enset sample, all analyses were performed in triplicate, and the base peak areas for detected amino acid derivatives analyzed using Thermo Xcalibur (Thermo Fisher Scientific, USA) and R software (R Core Team, 2017), and averaged per sample.

### 2.4 Genomic characterization of enset fermentation

DNA was extracted from kocho using a standard CTAB protocol. High quality DNA was successfully isolated from the *kocho* of three landraces (Ganticho, Agade, and Ado), one sample failed (Hala). Library preparation followed the double digest restriction site associated sequencing protocol (tGBS) of Data2Bio (Iowa, USA). Sequencing was performed using the Ion Proton platform (Thermo Fisher Scientific, USA). Raw reads were subsequently trimmed and quality controlled following the Data2Bio pipeline.

To identify microorganisms present in *kocho* samples, retained sequences were mapped to 24,610 reference bacteria and 285 reference fungi genomes, obtained from NCBI (https://www.ncbi.nlm.nih.gov/, accessed 20th March 2019). A local blast database was built and blast searches performed with e-value threshold of e^−2^ and scoring (match-mismatch) =1-2. The percentage of retained reads that aligned to a bacterial or fungal genome were recorded. As a control, DNA was also extracted from the leaf tissue of multiple farmer-grown enset accessions and a similar pipeline used to characterize bacteria or fungi present on the surface or within plant tissue. This approach provides a comparative line of evidence that where specific microbes identified in kocho samples are present at high concentrations, this is indicative that they are associated with fermentation (rather than simply present). It also provides an indication on whether the microbial community responsible for fermentation originates from the plant surface/tissue or elsewhere, such as the wider environment (e.g. soil) or a starter culture.

## 3. Results and discussion

### 3.1 Sample collection, processing and associated knowledge

Fermentation was initiated from mashed pseudostem tissue left for several days at ambient temperature without contrived additives of cultures or previous products. Kocho and Bulla were then fermented for three months, and Ko-Bu was fermented for six months. We recorded the ubiquitous use of fresh enset leaves as a work surface for processing and the prevailing views that fermentation will not be successful if the fermentation pit is not positioned within the enset growing area. Positioning of the enset fermentation pit within the enset growing area may indicate that temperature (or shade) is important, or alternatively may be related to the presence of specific microorganisms in the soil or elsewhere in the environment associated with enset plants. The resulting fermentation products can be stored for long periods (>6 months). As expected, farmers reported that different enset landraces produce *kocho* (or other products) of different qualities or preferred uses (including as medicines), however multiple landraces are normally processed together. In times of extreme famine, wild enset may be blended with harvested domestic plants, despite generally being considered unpalatable, though the resulting product is regarded as having lower quality, perhaps due to taste.

### 3.2 Inorganic composition of enset and comparison with other regional crops

Mean ash content across 19 corm samples was 5.0% (sd=1.4%), which is similar to previously published ash content (4.5%) for enset corms (Nurfeta et al., 2008). Concentrations of inorganic micronutrients and trace elements in raw enset corm tissue were averaged across 19 corm samples, each with two replicates, and plotted in Figure 1A. Inorganic micronutrient composition varied both by individual sample, and region (Figure 1B), although these data are unable to distinguish the relative importance of these two factors. Samples originating from Gurage zone were most variable on the first axis (27.1% of the variation), whilst samples from Sidama vary on the second axis (22.7% of the variation).

**Figure 1.**
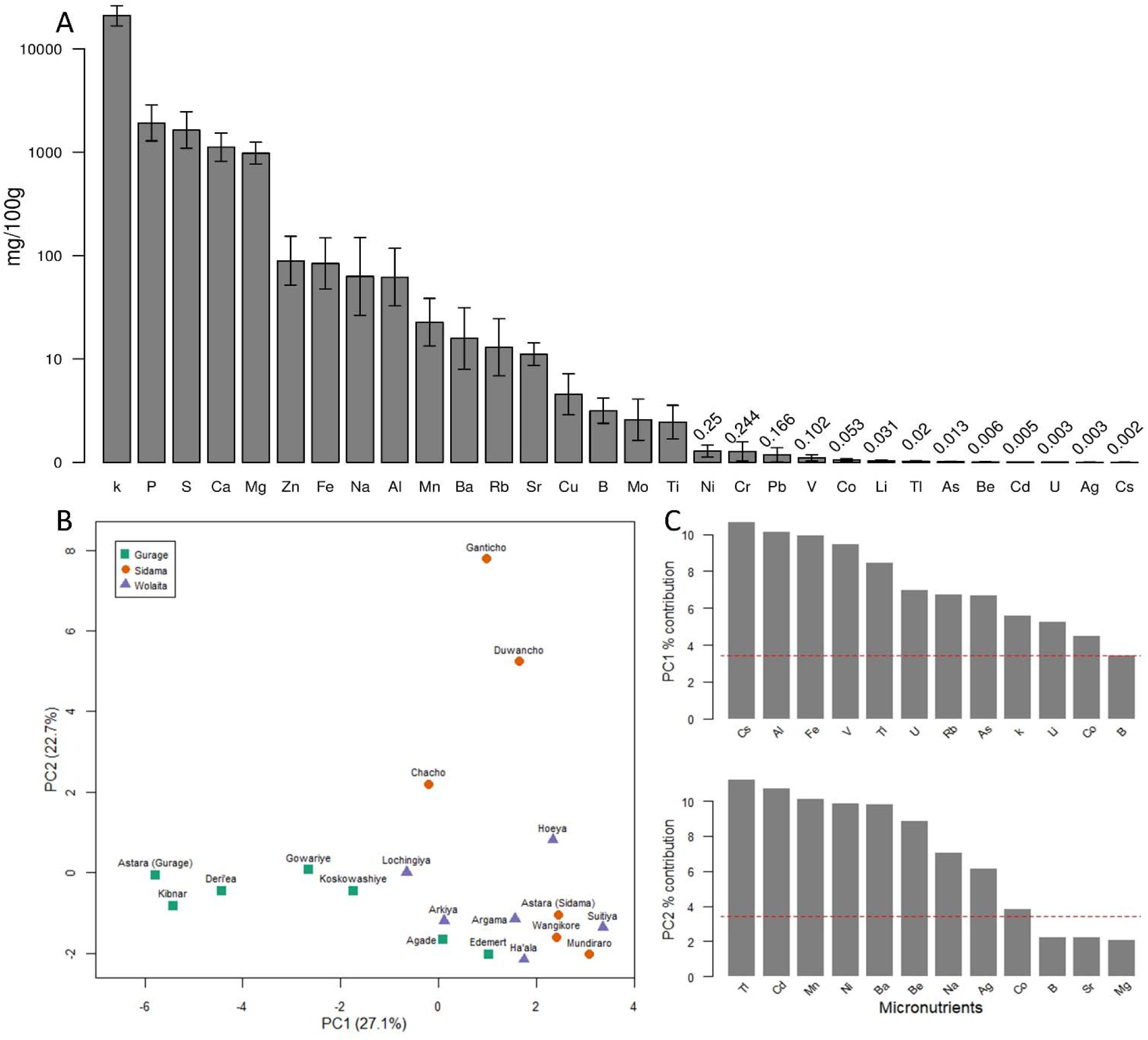
Concentration of inorganic minerals in enset corm tissue. A) Mineral values averaged across 19 enset corm samples. Bars denote standard deviation. B) Principal component analysis of enset corm inorganic composition across three geographical regions. C) Axis loading plots for the first and second principal components. The horizontal red line shows the level at which axis contributions would be equal.

These data illustrate important variation in micronutrient composition across enset landraces and growing regions, which mirrors the high vernacular and genetic diversity reported elsewhere for this crop (Borrell et al., 2018; Yemataw, Mohamed, Diro, Addis, & Blomme, 2014). This indicates the influence of genetic, environmental and/or management factors, and as a result, there are likely to be appropriate genetic targets for breeding to enhance nutritional composition. Similar performance gains have been achieved in a range of other species for which there is high variability in micronutrient composition (Welch & Graham, 2004). These data also suggest that climate or soil conditions, together with enset management practices may influence the uptake and accumulation of micronutrients in enset corm tissue, in particular because micronutrient composition appears to cluster weakly by region (Figure 2) – though many landraces are region specific making further investigation of this pattern challenging. We note that consumption of some micronutrients in high amounts is not recommended (e.g. Ni, Cd) and high levels of these metals in samples from Sidama (Figure 1B, 1C) may be associated with groundwater contamination.

**Figure 2.**
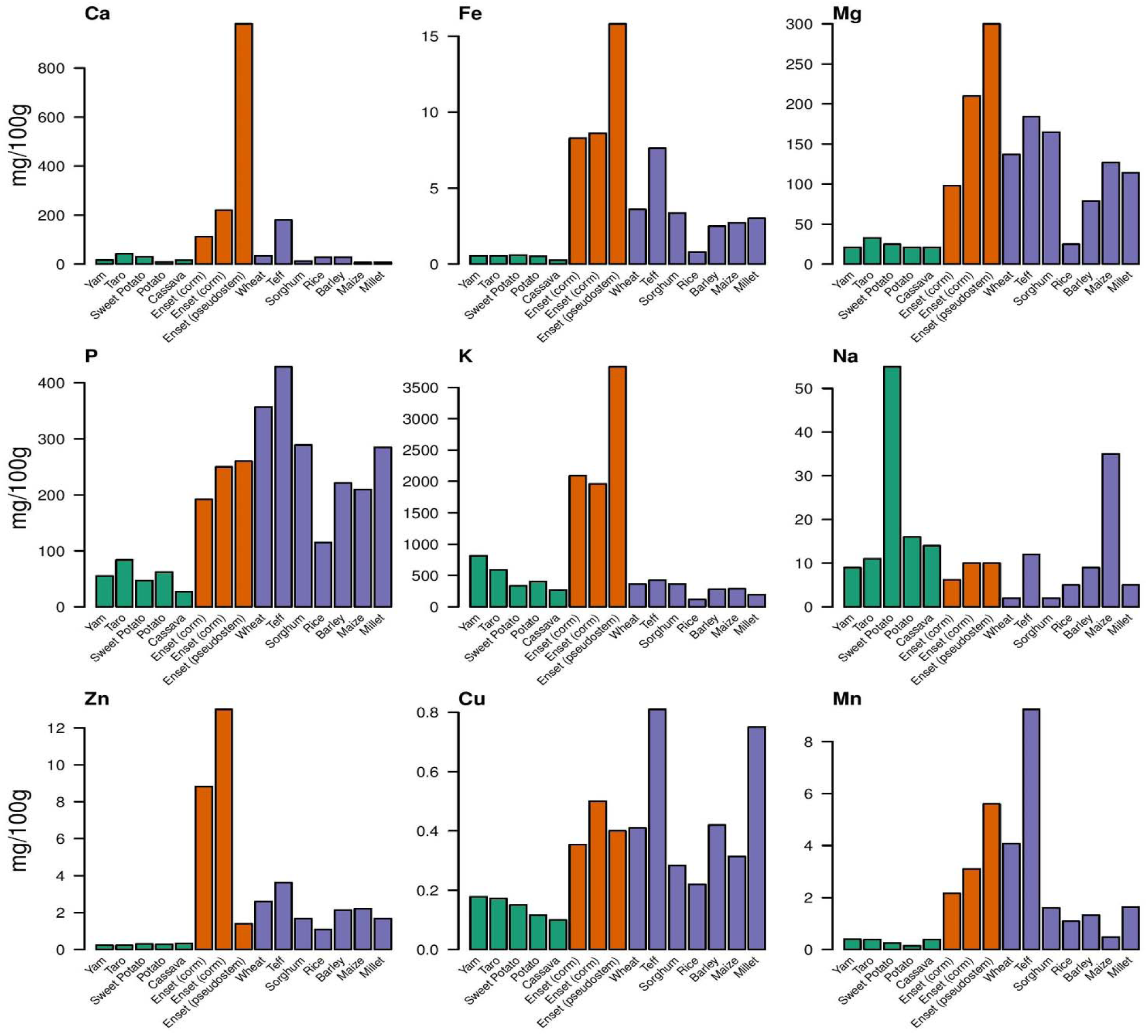
Comparison of enset tissue inorganic micronutrients (corm and pseudostem) with regionally occurring tubers and cereal crops. For enset three values are provided (from left to right); i) Enset (corm) results from this study (note: unavailable for K), ii) Enset (corm) from published sources and iii) Enset (pseudostem) from published sources. Comparative values for tuber and cereal crops are sourced from Feedipedia and the USDA Food Composition Databases.

In a comparison with regionally important tuber and cereal crops, enset reports particularly high values for calcium, iron, potassium and zinc and relatively low values for sodium (Figure 2). Both iron and zinc deficiencies are widespread in the region (Grebmer et al., 2018). Anemia (iron deficiency) is reported in 56% of children and 24% of adult women in Ethiopia (Gebru et al., 2018), with anemia also included in the World Health Organization’s Global Nutrition Monitoring Framework for Ethiopia (WHO, 2019). This emphasizes the importance of enset as a potential dietary source of iron. Iron concentrations have previously been reported as higher in enset pseudostem compared to corm (Heuze et al., 2017) and we find further support for this pattern across a larger range of landraces (Figure 2), though we show almost 8-fold variation between samples. Zinc deficiency is also reported in diets in Ethiopia (Gebru et al., 2018; Kebede & Modes, 2013). In this study we provide further evidence that enset is an important source of Zinc, and that enset corm contains higher levels of zinc than pseudostem tissue. Overall, most micronutrients occur at higher concentrations in the pseudostem than the corm (Figure 2).

### 3.3 Organic composition and the nutritional implications of enset fermentation

Mean total protein content across 19 corm samples in this study was 4.6% (sd=1.5%), broadly consistent with values published elsewhere (Feedipedia: 3.5%, sd=1.1%). The mean concentration of FAAs across raw enset tissues and processed food products are reported in Figure 3A with overall concentrations for each FAA shown in Figure 3B. Of the edible components of enset, corm tissue contained the highest concentrations of FAAs, but was also most variable. FAA variation across samples is presented in Figure 3C with variable loadings for the first and second principal components reported in Figure 3D. Based on principal component analysis, concentrations for pseudostem were mostly clustered but showed higher variation for corm tissue. The most variable FAAs on the second axis associated with corm variability included arginine, histidine and aspartic acid. The relative concentrations of individual FAAs before and after fermentation is reported in Figure 4.

**Figure 3.**
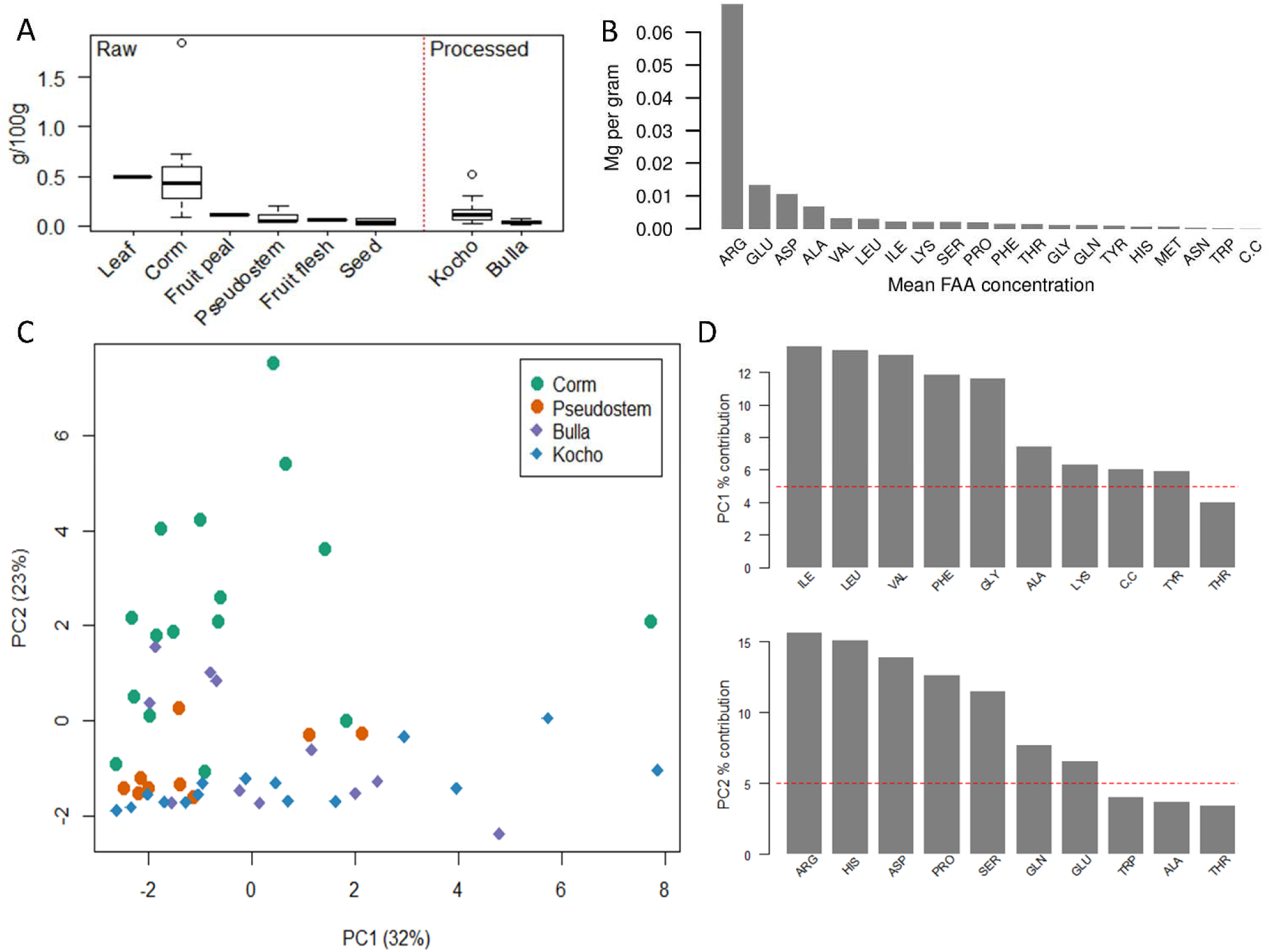
Free Amino Acid composition and variation in enset. A) Quantities of Free Amino Acids (FAAs) present across raw enset tissues and processed enset food products. B) Mean concentrations of free amino acids across all samples. C) Principal component analysis of enset FAAs across four tissue types. D) Axis loading plots for the first and second principal components.

**Figure 4.**
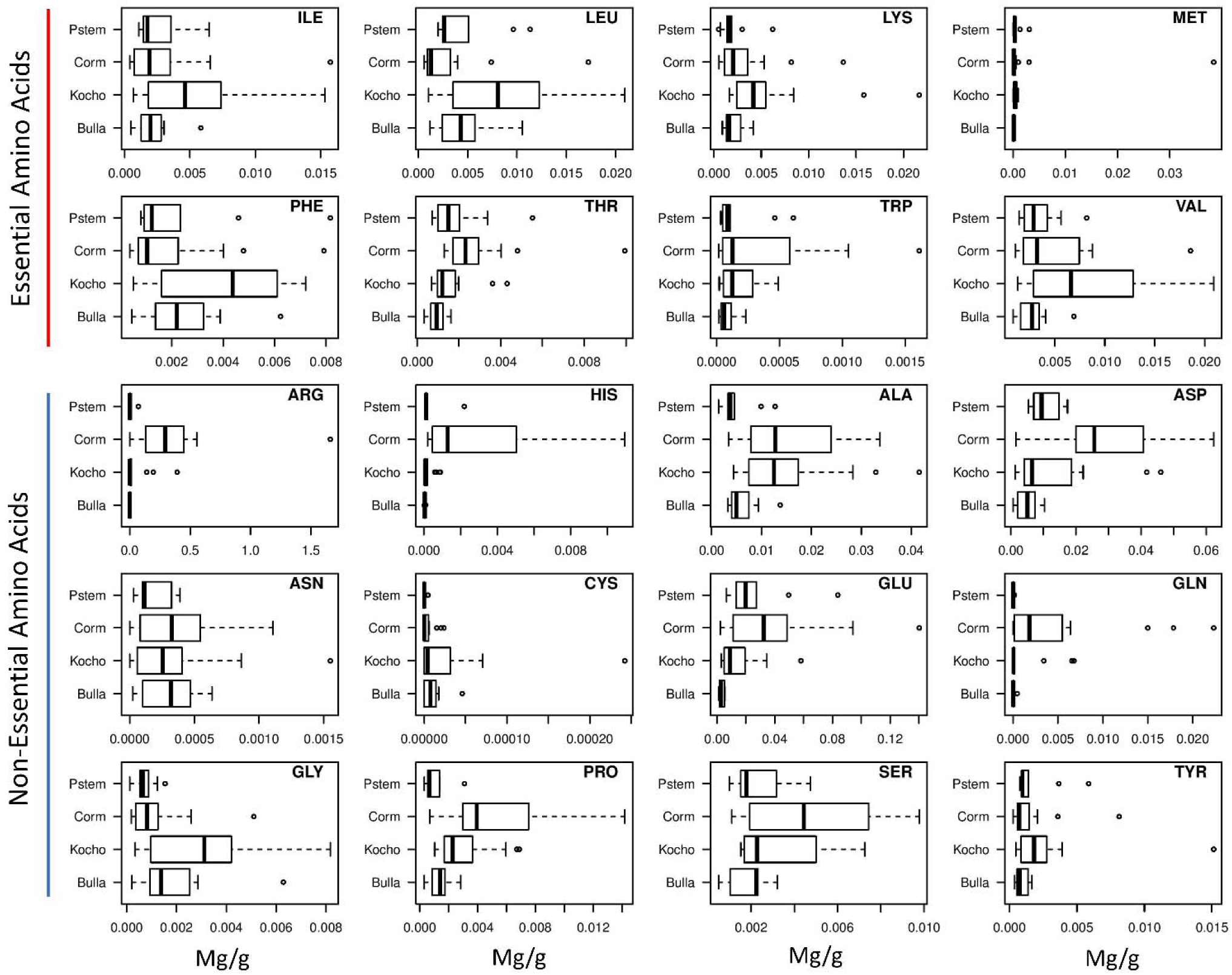
Free Amino Acid composition in raw enset tissues (pseudostem and corm) and processed food products (*kocho* and *bulla*). Whilst essential and non-essential amino acids are noted, conditionally essential amino acids, required in certain circumstances, are presented and include arginine, histidine, glycine and glutamine.

Processing of crops is normally performed to improve the digestibility of the product, removing anti-nutritionals and allowing storage without microbial or animal (including insect) contamination. However, there is no evidence that unprocessed enset is toxic, contrasting with cassava (cyanide) or many legumes (with both highly toxic lectin glycoproteins and frequent association with fungi producing mycotoxins). In this instance, enset processing appears to have a significant impact on the composition of derived food products, with FAAs present at higher concentrations in raw tissues (particularly leaf and corm tissue; Figure 4A), suggesting that microorganisms are net consumers of FAAs during enset fermentation. These data also show that certain FAAs may be localized to specific tissues, for example arginine and glutamine are both detected in corm, but at low concentrations or absent in other tissues both pre- and post-fermentation (Figure 4). Similarly, phenylalanine and glycine occur at higher concentrations in fermented products than raw tissues, whilst many other FAAs decline. Overall, several essential amino acids (isoleucine, leucine, phenylalanine and valine) are increased in fermented products, particularly the main staple food, *Kocho*, suggesting that traditional fermentation practices may contribute to nutrition.

When protein or amino acids are ingested, the vast majority of digestion products that reach the blood stream are single amino acids, with the completeness of protein digestion dependent on metabolic factors (Bhutta & Sadiq, 2012). Thus in some cases ingested FAAs may be more bioavailable whilst in others a protein-rich diet may be poorly digested and of reduced nutritional value (Bhutta & Sadiq, 2012). Furthermore, certain amino acids contribute to the flavour of foods (Kato, Rhue, & Nishimura, 1989), thus influencing the selection of processing or fermentation methods and dietary choices of consumers, with potential consequences for human nutrition. We note wild enset is not consumed because it is considered bitter and unpalatable. It is interesting, therefore, that fermentation appears to markedly reduce aspartic acid and glutamic acid, both of which produce a potent sour taste at relatively low concentrations, as well as reducing arginine and histidine concentrations, which are characteristic of bitter tastes. Whether development of processing techniques has been concomitant with domestication to produce food with improved palatability is an interesting area for further research.

This study also reveals detection of varying concentrations of the essential amino acids isoleucine, leucine, lysine, phenylalanine, threonine, tryptophan, and valine in different enset tissues (Figure 4), in the form of FAAs. Consequently, these data provide the first evidence that enset, as part of a broader diet, may contribute to intake of these essential amino acids. The principal FAA detected across all enset samples analysed in this study was arginine (Figure 4), which occurred at the highest levels in corm tissue. Although arginine is not an essential amino acid, some evidence suggests that increased dietary arginine can improve outcomes in critically ill individuals (Emery, 2012) and it is considered essential for infant growth, with histidine also being important for the latter (Brayfield, 2019), and also detected in enset tissue, especially the corm (Figure 5).

**Figure 5.**
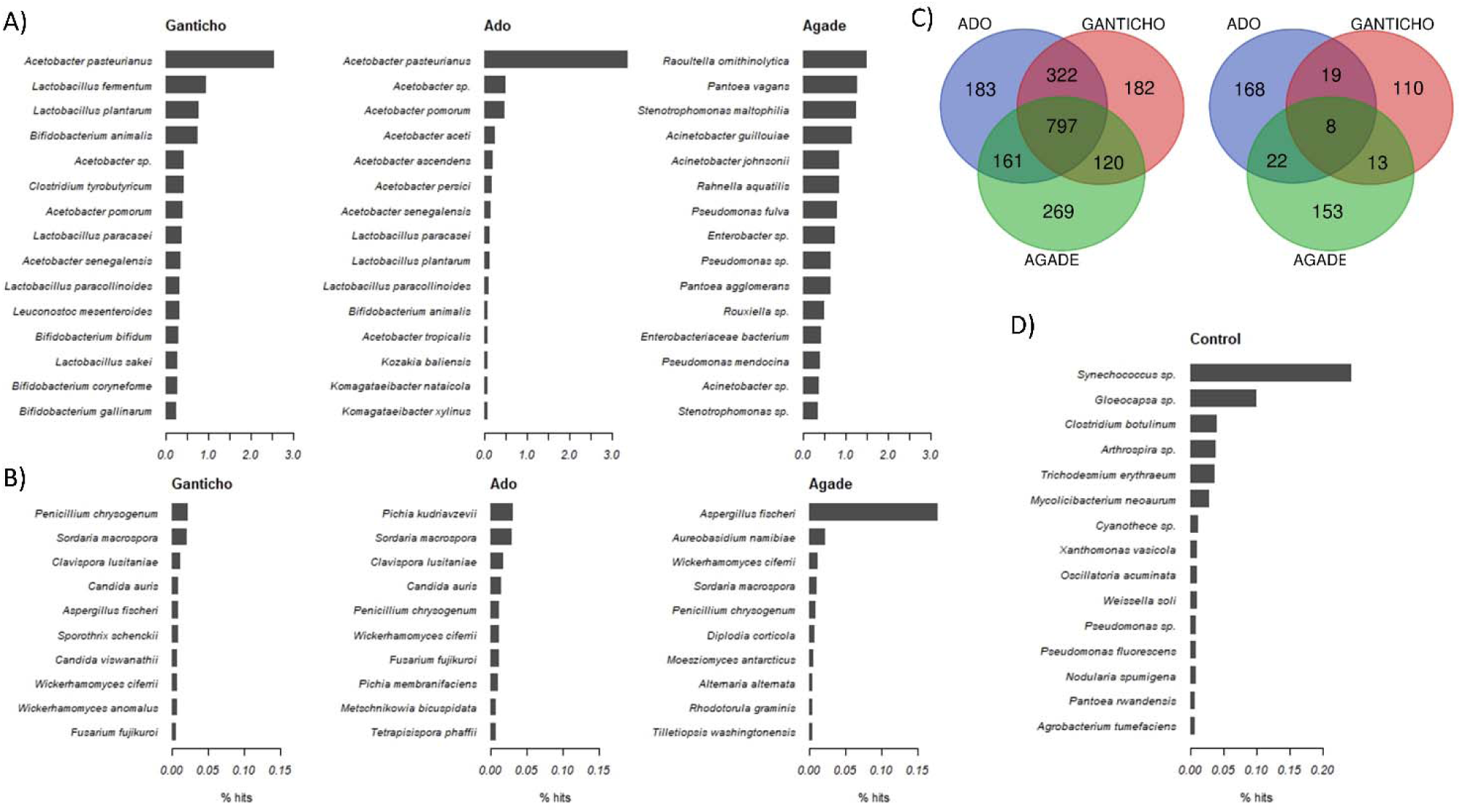
Microbial composition of enset kocho samples. Bacteria (A) and Fungi (B) genomes with highest percentage hit rates across three kocho samples. C) Venn diagrams illustrating the number of species in common between samples for Bacteria (left) and Fungi (right). D) Top bacteria and fungi hits from enset leaf tissue

A traditional use of enset in Ethiopia is its intake in the form of amicho (boiled corm), which is reputed to heal bone fractures (Borrell et al., 2018). Arginine is involved in collagen formation, tissue repair and wound healing via proline, which is hydroxylated to form hydroxyproline, and it may also stimulate collagen synthesis as a precursor of nitric oxide (Van de Poll, Luiking, Dejong, & Soeters, 2012). The analysis of free amino acids in this study has revealed high levels of arginine, compared to other amino acids, detected in different edible parts of enset. This novel finding provides a rational scientific basis for the first time that may explain the reputed traditional use of enset to aid bone healing after breakage. Intriguingly, three landraces often reported as having medicinal properties (Koshkowashiye (Gurage), Astara (Sidama) and Lochingiya (Wolaita)) report three of the four highest arginine values, whilst Lochingiya and Astara also report the highest calcium concentrations, a mineral critical for bone development if not healing.

### 3.4 Microbial community characterization and signatures of contamination

Sequencing of kocho samples resulted in 9.6M raw reads. After quality control 8.6M reads were retained with a length of 50-209bp. Of these, 1.9M and 24,300 reads returned BLAST hits above a threshold of e^−2^ for bacteria and fungi genomes, respectively. The bacterial species *Acetobacter pasteurianus* was the most frequent in the *kocho* samples Ganticho and Ado, whereas *Raoultella ornithinolytica* was found to be most frequent in Agade (Figure 5A). The fungal species *Penicillium chrysogenum*, *Pichia kudriavzevii* and *Aspergillus fischeri* were found most frequent in the Ganticho, Ado and Agade samples respectively (Figure 5B). A total of 797 bacterial species were found in common across the three kocho samples, though comparatively only eight fungi species were found across all samples (Figure 5C). As a control, BLAST results from leaf tissue is provided in Figure 5D and largely identifies distinct species from the main kocho analysis. The same analysis is presented at genus level in Figure S1.

Genomic analysis of the microbial community associated with enset fermentation showed that the most abundant genus of bacteria was *Acetobacter*, which is generally aerobic and known for producing acetic acid, reducing the pH of the fermenting enset pulp. It both gives a desirable flavor to the product and lowers the pH which inhibits growth of other organisms, thus allowing safe storage of the product. Whilst these species may occur as endophytes, the most common species identified in the kocho samples (*Acetobacter pasteurianus*) was not identified in raw leaf tissue (Figure 5D). The second most abundant genus was *Lactobacillus*, a group of anaerobic bacteria often associated with controlled fermentation in foods. In a comparison with the bacteria identified by Gashe (1987) we also found *L. mensenteroides*, *L. coryniformis* and *L. plantarum* but not *S. faecalis*. In the genome sequencing of leaf-derived DNA, Harrison et al., (2014) reported extensive hits (>8%) against *Pseudomonas fluorescens* and *Methylobacterium radiotolerans*, which they propose are endophytes associated with enset; we also find a small number of *P. fluorescens* hits in our kocho samples, but only one hit to *M. radiotolerans*. Overall, our analysis identified many more bacterial than fungal sequences. We also find a much higher proportion of bacteria species in common between samples (39.2%), compared to fungi (1.6%) including the yeasts. This suggests that bacteria are principally responsible for enset fermentation.

These analyses provide an opportunity to identify potentially harmful microorganisms such as those associated with spoilage or food poisoning (Bhunia, 2018). The number of hits associated with *Escherichia, Campylobacter, Salmonella, Clostridium* (except in the *Ganticho* kocho sample) and Listeria was generally very low in all samples. However, whilst the microbial community composition of Ganticho and Ado were very similar, Agade was dominated by *Raoultella ornithinolytica*, a species associated with human infections. *Stenotrophomonas maltophilia*, *Acinetobacter johnsonii*, *Rahnella aquatilis* are other species potentially harmful to health were also among the most frequently identified bacteria in this sample. Similarly, the highest fungal hit for Agade was *Aspergillus (Neosartorya) fischeri*, a close relative of the major pathogen *Aspergillus fumigatus* also associated with hypersensitivity pneumonitis (farmer’s lung, immunologically mediated inflammation). This suggests that in comparison to Ganticho and Ado, Agade could be characterized as a contaminated food product. The prevalence of enset food product contamination in Ethiopia is unknown.

## 4. Conclusions

Ethiopia has historically been the world’s largest recipient of targeted food aid (World Food Programme, 2013) and is 93^rd^ of 119 qualifying countries in the 2018 Global Hunger Index (Grebmer et al., 2018). Nationally, undernourishment affects 21.4% of the population and 38.4% of children under the age of five are affected by stunting (Grebmer et al., 2018). Therefore effective utilization of agricultural diversity is a priority to achieve food security and address public health needs, particularly in the context of climate change and population growth (Pironon et al., 2019). However despite being a center of diversity for plant domestication, dietary diversity over much of Ethiopia is extremely low due to overdependence on starchy staples (Gebru et al., 2018) with major dietary deficiencies in iron and zinc. This is indicative of a However, whilst child stunting prevalence in the enset growing region (reflecting chronic undernutrition) is largely consistent with the national average (CSA and ICF, 2016), these areas have some of the lowest national levels of child wasting (an indicator of acute undernutrition). This provides compelling indirect evidence of the food security potential of enset in mitigating acute food insecurity events.

The results presented here show that there is significant potential for enhanced nutritional benefits (e.g. iron, zinc, FAAs) from enset that could impact the chronic health and welfare challenges experienced by millions of Ethiopian farmers whom rely on enset as a resilient starch staple. We show that there is significant variation in enset nutritional diversity, partitioned across multiple stages of enset cultivation and processing from the selection of landraces, environmental conditions and management practices, to the timing and selection of tissues for harvest and the microbial community associated with enset processing. Many of these sources of variation are not currently understood, controlled or investigated and represent significant opportunities for optimization or improvement. We also highlight that in addition to there being few enset germplasm collections (Borrell et al., 2018), there are no collections of the microbial communities associated with kocho processing. We therefore suggest that it is important to further document, collect and preserve their diversity as they would be lost should farm structures, agronomy or management change for social or economic reasons. In summary, more than 20 million Ethiopians rely on enset-derived products as a starch staple or co-staple, and this population is projected to grow significantly in the coming decades (Borrell et al., 2018). Therefore, selection of enset landraces with improved raw nutritional content or enhanced processing techniques that improve the composition, quality or safety of enset based foods has the potential for significant public health impacts.

## Acknowledgements

We thank farmers for processing enset tissues for study, and field assistants for data collection.

## Ethics Statement

All indigenous knowledge associated with enset was collected with prior informed consent and in accordance relevant Access and Benefit Sharing Agreements.

