## Supplementary Figure 1 for "Micronutrient composition and microbial community analysis across diverse landraces of the Ethiopian orphan crop enset"

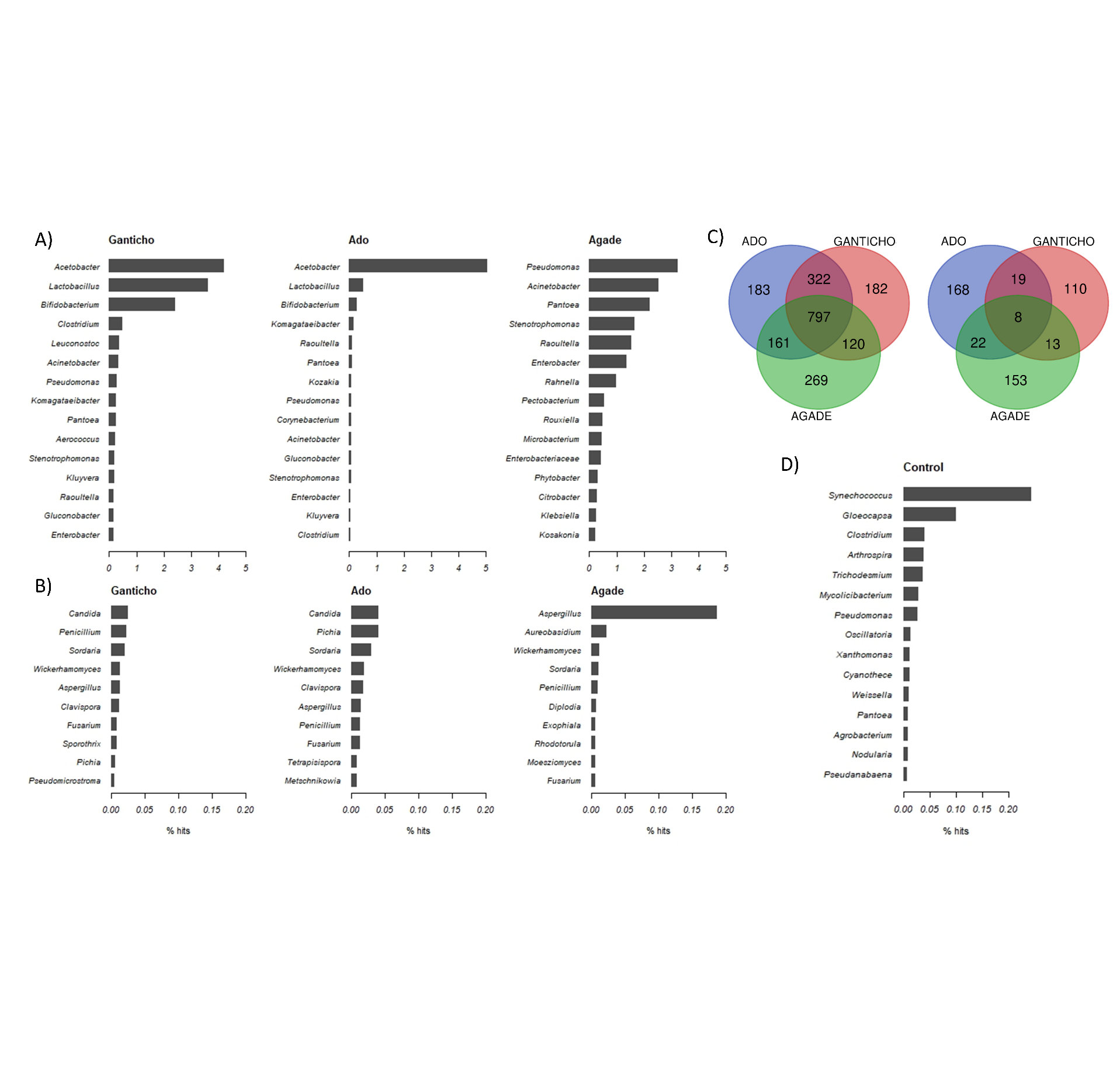

**Figure S1.** **Microbial composition of enset kocho samples AT GENUS LEVEL.** Bacteria (A) and Fungi (B) genera with highest percentage hit rates across three kocho samples. C) Venn diagrams illustrating the number of genera in common between samples for Bacteria (left) and Fungi (right). D) Top bacteria and fungal genera hits from enset leaf tissue.
